# Prenatal THC exposure disrupts mitochondrial respiratory gene programs and medium spiny neuron maturation trajectories in the nucleus accumbens

**DOI:** 10.64898/2026.04.26.720961

**Authors:** Zhong Chen, Wanqiu Chen, Yun Seok Lee, Wendell Jones, Laura Goetzl, Jennifer Dawn Thomas, Yan Dong, Charles Wang

## Abstract

Prenatal cannabis exposure (PCE) is increasingly prevalent and has been associated with adverse neurodevelopmental outcomes, yet its molecular impact on brain reward circuitry remains poorly defined. Here, we investigated transcriptional and epigenomic alterations in the nucleus accumbens (NAc) following prenatal Δ^9^-tetrahydrocannabinol exposure in a rat model using snRNA-seq and snATAC-seq analyses. PCE markedly suppressed the expression of genes involved in mitochondrial oxidative phosphorylation (OXPHOS) in the NAc on postnatal day 24 (P24), indicating reduced mitochondrial respiration capacity. This disrupted mitochondrial respiratory gene programming was accompanied by coordinated alterations in ribosomal and proteasomal pathways regulating protein homeostasis in NAc medium spiny neurons (MSNs), suggesting coupled disruption of cellular metabolism and neuronal maturation. snATAC-seq analysis revealed altered chromatin accessibility at promoter regions enriched for *Nrf1* and *Yy2* binding motifs, implicating *Hrf1-* and *Yy2-*associated transcriptional regulation of mitochondrial genes in MSNs following PCE. Moreover, an acute THC challenge in PCE offspring at P24 further exacerbated the suppression of genes involved in mitochondrial OXPHOS and MSN maturation. Together, these findings define a transcriptional and epigenetic framework through which PCE may perturb mitochondrial function and impair MSN maturation trajectories in the NAc, providing mechanistic insights into how PCE may alter the development of reward circuitry.

## Introduction

Cannabis is one of the most commonly used substances in pregnant women^1,2^. A recent study reported a prevalence as high as 22.4% for detectable prenatal marijuana exposure based on umbilical cord blood testing^3^. Prenatal cannabis exposure (PCE) poses health risks to the developing fetus because 7^9^-tetrahydrocannabinol (THC), the primary psychoactive constituent of cannabis, readily crosses the placenta to disrupt the endogenous cannabinoid system, leading to long-term neurodevelopmental disorders including impaired cognitive function, hyperactivity, and increased impulsivity^4–6^. Given the rising prevalence of maternal cannabis use and its deleterious effects, it is critical to investigate the impact of PCE on fetal brain development.

The nucleus accumbens (NAc) is a key brain region regulating emotional and motivational responses to both natural rewards and drugs of abuse^7–9^. Cannabis exposure has been associated with alterations in the neural architecture of core reward structures^10^. For example, animal studies have shown that cannabinoids induce structural abnormalities^11^ and altered synaptic transmission in the NAc^12^. However, it remains unclear how PCE affects the development and function of the NAc at cellular and molecular levels.

Mitochondria undergo morphological changes and a metabolic shift from glycolysis to oxidative phosphorylation during differentiation in both cardiac and motor neurons^13,14^. Similar changes, including increased mitochondrial oxygen consumption, have been observed during embryonic stem cell (ESC) differentiation^15^. Inhibition of mitochondrial respiration impairs cellular differentiation and promotes maintenance of stem cell pluripotency^16^, while mitochondrial oxidative activity and glucose metabolism play a key role in regulating the tempo of neuronal maturation^17^. Emerging evidence further suggests that mitochondrial metabolism not only supports neuronal energy demands but also regulates transcriptional and epigenetic programs that govern neuronal differentiation and maturation.

Biochemical and cytological studies indicate that THC disrupts mitochondrial function in the brain^18^. Several molecular mechanisms have been proposed. Initially, THC was thought to act directly on mitochondria due to their high lipophilic membranes, potentially disturbing mitochondrial membrane properties via destabilization of cardiolipin^19^. Subsequent studies suggest that THC can also regulate mitochondrial calcium levels, impair the electron transport chain, disrupt oxidative phosphorylation (OXPHOS) and ATP production, or alter mitochondrial biogenesis dynamics^18,20^. Collectively, these findings suggest that THC-induced mitochondrial dysfunction may contribute to the adverse developmental effects of PCE. However, the temporal and cell-type specific transcriptional consequence of PCE within circuitry remain poorly understood.

We hypothesized that PCE disrupts the expression of nuclear-encoded mitochondrial genes in NAc neurons through altered transcriptional regulation and chromatin accessibility, thereby perturbing neuronal maturation programs. To test this, we performed snRNA-seq and snATAC-seq on NAc tissues from postnatal day 24 (P24) PCE rats, along with snRNA-seq following acute THC challenge in PCE male offspring. PCE suppressed the expression of genes involved in the mitochondrial oxidative phosphorylation, with the most pronounced effects observed in medium spiny neurons (MSNs). These transcriptional alterations were accompanied by coordinated changes in ribosomal and proteasomal pathways, suggesting disruption of cellular metabolic and maturation programs. Acute THC challenge in PCE offspring further exacerbated the suppression of mitochondrial respiratory genes and altered MSN maturation-associated transcriptional signatures. Together, these findings suggest that PCE may increase the vulnerability of developing reward circuitry to subsequent cannabinoid exposure.

## Results

### PCE induces transcriptomic and epigenetic changes in rat NAc

We simultaneously profiled gene expression (GEX) and chromatin accessibility (ATAC-seq) from the same single nuclei isolated from the NAc, using the 10x Multiome chemistry. The nuclei were isolated from a pool of NAc tissues collected from four P24 rats derived from three dams across four groups: female control (F_Ctl), female prenatal cannabis exposure (F_PCE), male control (M_Ctl), and male prenatal cannabis exposure (M_PCE). Prenatal THC exposure was administered from embryonic day 5 (E5) to E20, spanning implantation through late gestation, a developmental window roughly corresponding to the late first trimester through term in humans. We obtained 24,095 nuclei with matched transcriptomic and chromatin accessibility profiles (F_Ctl: 6,203; F_PCE: 6,446; M_Ctl: 4,838; M_PCE: 6,608) after integrating the GEX and ATAC-seq data and performing quality control based on RNA and ATAC features (nFeature and nCount). Unsupervised dimensionality reduction using weighted nearest neighbors (WNN) identified 18 transcriptionally distinct cell populations (**Fig. 1a),** which were well mixed across groups **(Suppl. Fig. 1a**). No substantial differences in RNA or ATAC quality metrics were observed among groups within clusters (**Suppl. Fig. 1b-c**), supporting the absence of technical bias in downstream analyses.

**Fig. 1.**
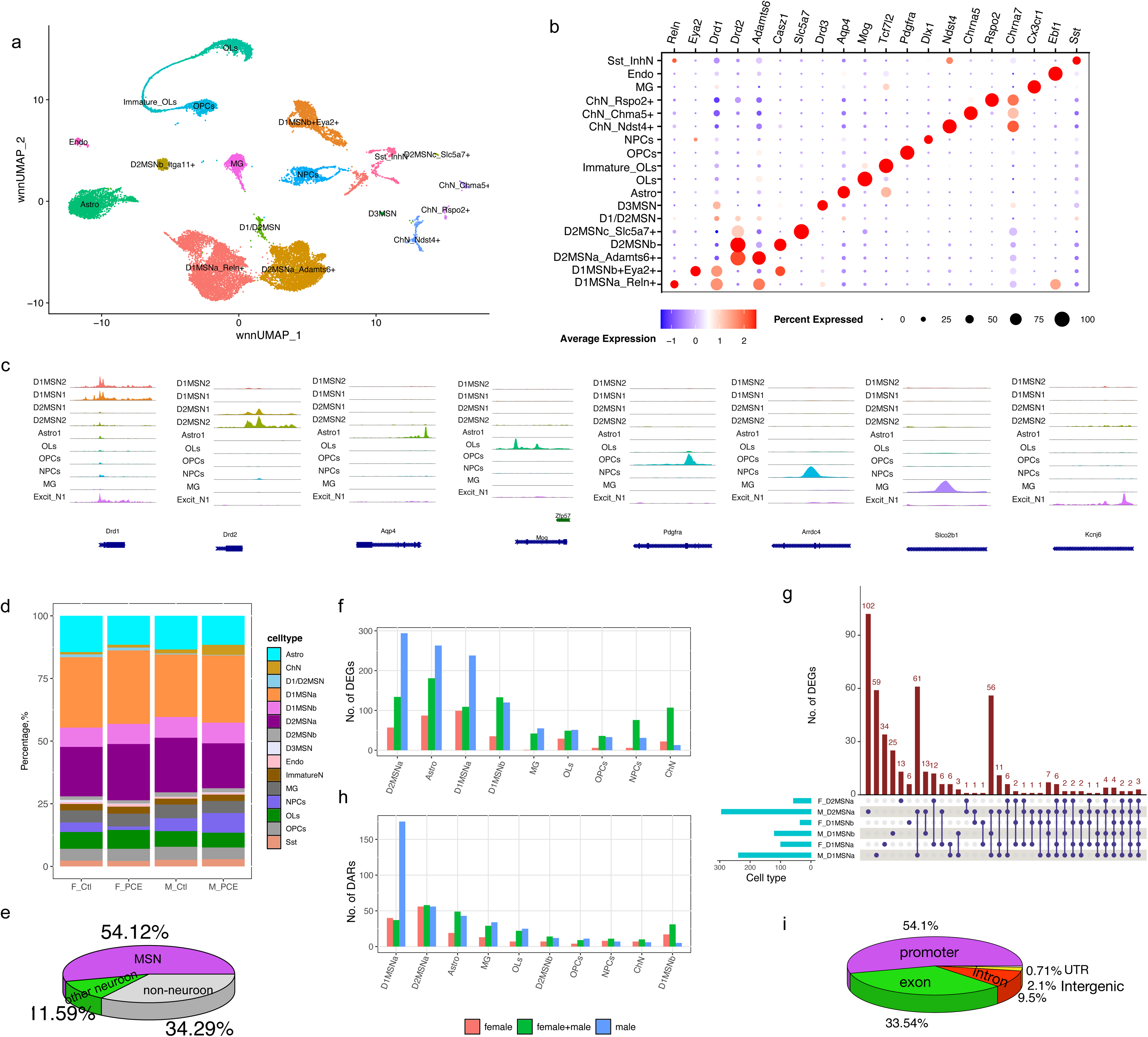
PCE induces transcriptomic and chromatin structural changes in P24 rat NAc. (**a**) Unsupervised cell clustering based on wnn_UMAP of Multiome data from F_Ctl, F_PCE, M_Ctl, and M_PCE groups. (**b**) Dot Plot showing the expression of cluster-specific marker genes. (**c**) Peak coverage tracks highlighting cell type-specific ATAC-seq signals across major cell populations. The three ChN subtypes were combined to increase ATAC signal intensity. (**d**) Bar plot illustrating cell-type composition across four datasets; clusters containing fewer than 20 cells were excluded. (**e**) Pie chart showing the proportion of all MSNs among all cells combined from four datasets. (**f, h**) Bar plots showing the number of DEGs (**f**) and DARs (**h**) identified in each cell type. (**g**) UpSet plot illustrating shared and unique DEGs among D1MSNa, D1MSNb, and D2MSNa populations in male and female PCE rat NAc. (**i**) Pie chart showing the genomic annotation distribution of differential peaks induced by PCE.

Cell type annotation based on canonical marker genes identified multiple neuronal and glial populations (**Fig. 1b**). Among medium spiny neurons (MSNs), we resolved two *Drd1*⁺ subtypes—D1MSNa (*Reln^+^*) and D1MSNb (*Eya2^+^*); three types of *Drd2*^+^ MSN: D2MSNa (*Adamts6^+^*), D2MSNb (*Itga11^+^*), and D2MSNc (*Slc5a7^+^*); as well as a *Drd3^+^* MSN population (D3MSN). Additional cell types included astrocytes (*Aqp4*), oligodendrocytes (OLs) (*Mog*), oligodendrocyte progenitor cells (OPCs) *(Pdgfra*), microglia (MG) (*Cx3cr1*), neuron progenitor cells (NPCs) (*Nol4*). A small population of cholinergic neurons with three different subtypes was also identified based on expression of the α-7 nicotinic acetylcholine receptor (**Fig.1b**).

Integration of chromatin accessibility with gene expression revealed concordance between accessible regulatory regions and marker gene expression across cell types (**Fig. 1b-c**). Analysis of cell-type composition showed no significant differences between groups following PCE (**Fig. 1d**). MSNs comprised 54% of all cells and over 80% of neurons (F_Ctl: 80.7%, F_PCE: 89.2%, M_Ctl: 82.3%, and M_PCE: 77.5%), consistent with prior reports of NAc cellular composition^21^ (**Fig. 1e**).

We next identified differentially expressed genes (DEGs) between PCE and control rats across major cell types (p_adj < 0.05 and |log2FoldChange| >0.35). PCE induced a greater number of DEGs in males than females across cell types (**Fig. 1f**). Within MSN subtypes (D1MSNa, D2MSNa, and D1MSNb), many DEGs were cell type specific (**Suppl. Table 1**). For example, 102 DEGs were unique to male D2MSNa, compared with 59 in male D1MSNa and 34 in female D1MSNa, highlighting cell-type- and sex-specific transcriptional responses to PCE (**Fig. 1g**). Few DEGs were shared between males and females within the same cell type, indicating pronounced sex-dependent effects. Conversely, a subset of DEGs was shared across multiple cell types (**Fig. 1g**), suggesting that PCE also perturbs certain common molecular pathways.

Chromatin accessibility changes were modest overall, with comparable numbers of differential accessibility regions (DARs) between sexes across most cell types. An exception was D1MSN, where males exhibited a greater number of DARs (**Fig. 1h**). Notably, MSNs displayed more extensive chromatin remodeling than other cell types, with DARs enriched in promoter regions (**Fig. 1i, Suppl. Table 2**), suggesting that PCE preferentially affect transcriptional regulation at gene initiation sites.

### PCE suppresses expression of genes involved in mitochondrial oxidative phosphorylation

Common DEGs induced by PCE in D1 and D2 MSNs were predominately associated with mitochondrial electron transport and ATP production (**Fig. 2a**). These genes were consistently downregulated in MSNs from both male and female offspring, with a greater magnitude of reduction observed in males (**Fig. 2b**). This coordinated downregulation of mitochondrial genes is an indicative of impaired oxidative phosphorylation (OXPHOS) and is consistent with mitochondrial dysfunction previously reported in PCE models^18,20^.

**Fig. 2.**
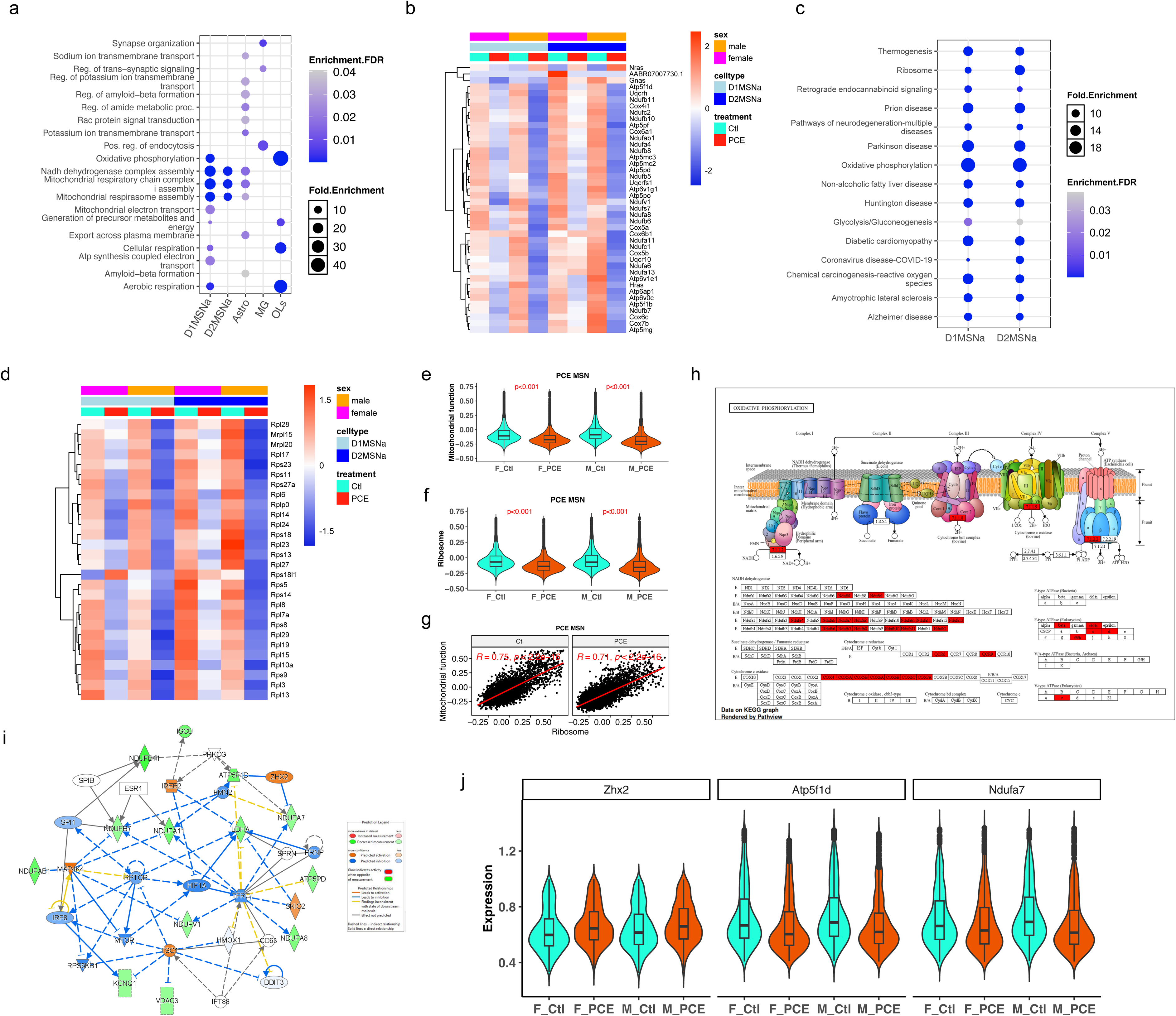
PCE induces mitochondrial dysfunction. (**a**) Dot plot showing the top enriched Gene Ontology biological processes (GO: BP) based on DEGs (combined male and female) induced by PCE across five major cell types. (**b**) Heatmap displaying the expression of D1MSNa and D2MSNa DEGs enriched in GO: BP. (**c**) Dot plot illustrating common KEGG pathways enriched from D1MSNa and D2MSNa DEGs. (**d**) Heatmap showing the expression of D1MSNa and D2MSN DEGs enriched in ribosomal assembly pathways. (**e, f**) Violin plots showing mitochondrial function scores (**e**), and ribosome module scores (**f**) in MSNs. (**g**) Pearson correlations between ribosome module scores and mitochondrial function scores in male MSNs, shown as a representative analysis. MSNs include combined D1MSNa and D2MSNa populations. (**h**) Schematic of mitochondrial oxidative phosphorylation (OXPHOS) with PCE-associated DEGs highlighted in red. (**i**) Ingenuity Pathway Analysis (IPA) network of OXPHOS enriched from combined male and female PCE DEGs. (**j**) Violin plots showing normalized expression levels of *Zhx2*, *Atp5f1d*, and *Ndufa7* at the single-nucleus level. For clarity, only D1MSNa and D2MSNa populations were shown, as they represent the predominant D1- and D2- MSN subtypes, respectively.

KEGG (Kyoto Encyclopedia of Genes and Genomes) enrichment analysis using DEGs from D1 and D2 MSNs further identified ribosome-related pathways as significantly affected by PCE (**Fig. 2c**). Correspondingly, ribosomal gene expression was broadly reduced, suggesting impaired translation capacity in NAc MSNs (**Fig. 2d**). Gene module scores based on mitochondrial and ribosomal pathway genes revealed coordinated decreases in both mitochondrial and ribosomal functional signatures (**Fig. 2e-f**). Notably, mitochondrial and ribosomal module scores were positively correlated (**Fig. 2g**), supporting a functional link between mitochondrial activity and protein translation which may be synergistically suppressed by PCE. These results suggest that suppression of ribosomal function may contribute to, or occur in parallel with, the perturbation of nucleus-encoded mitochondrial gene transcription in PCE-exposed neurons.

Both mitochondrial and ribosomal deficits were more pronounced in male MSNs compared to females (**Suppl**. **Fig. 2a, b**), indicating sex-dependent vulnerability. In males, additional DEGs were enriched in synaptic vesicle cycle and retrograde endocannabinoid signaling pathways (**Suppl. Fig. 2e**). This heightened sensitivity may be attributed to the increased expression of the cannabinoid receptor CB1 in males (**Suppl. Fig. 2f**), potentially amplifying the THC effects.

Beyond MSNs, similar reductions in mitochondrial and ribosomal gene module scores were observed in astrocytes (**Suppl. Fig. 2g, h**), although the correlation between these modules was weaker than in MSNs (**Suppl. Fig. 2i, g),** suggesting cell-type–specific coupling between metabolic and translational processes.

At the pathway level, downregulated mitochondrial genes were primarily associated with key OXPHOS components, including NADH dehydrogenase (Complex I), cytochrome c oxidoreductase assembly (Complex III/IV), and ATP synthesis (Complex V) (**Fig. 2h**). Upstream regulator analysis by IPA identified Zhx2, a Zinc-finger and homeoboxes 2 transcription factor previously implicated in metabolic regulations of mitochondrial function^22^, as a potential key mediator of PCE-induced effects on energy production (**Fig. 2i**). *Zhx2* transcription was increased in MSNs following PCE (log2FC=0.69), which was associated with reduced expression of its predicted target genes, including *ATP5f1d* and *Ndufa7* (**Fig. 2j**). This supports a model in which PCE disrupts OXPHOS through regulating the transcription of nuclear-encoded mitochondrial genes.

### PCE alters chromatin accessibility at genes involved in mitochondrial function and protein homeostasis

To investigate epigenetic mechanisms underlying PCE-induced transcriptional changes, we integrated gene expression with ATAC-seq data from male NAc cells, where PCE effects were most pronounced. In MSNs, we identified 389 DEGs and 476 genes associated with differential accessibility regions (DARs); however, only17 DEGs overlapped with genes harboring DARs, including 12 with promoter-associated changes (TSS±1k bp) (**Fig. 3a**). Among these genes, chromatin accessibility was strongly correlated with transcriptional output: 11 exhibited concordant decreases in promoter accessibility and gene expression,

**Fig. 3.**
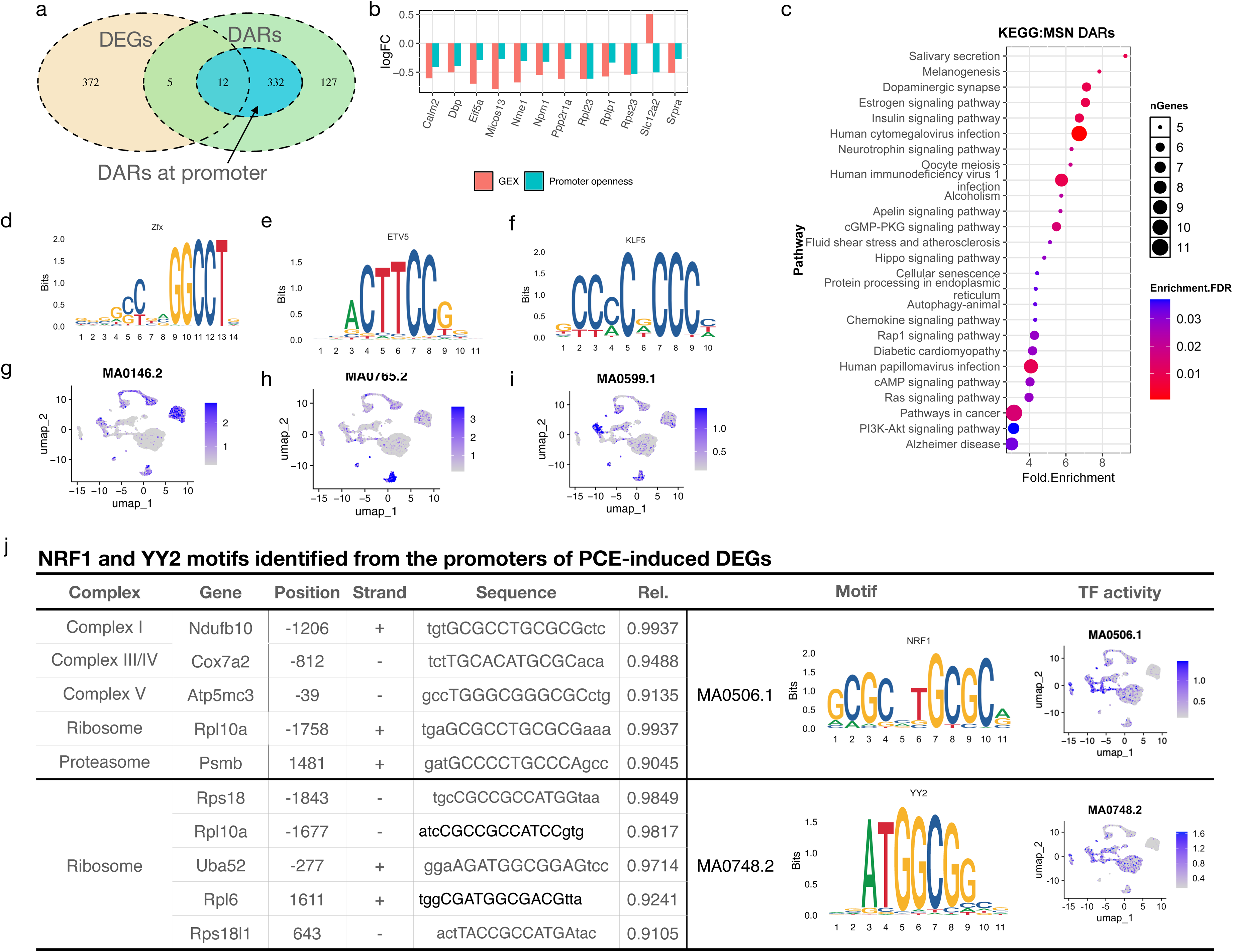
PCE alters the chromatin accessibility of mitochondrial and ribosomal genes. (**a**) Venn diagram showing the overlapped genes between PCE-induced unique DEGs and genes associated with DARs in D1 and D2 MSNs. (**b**) The Log2 fold change (log2FC) of gene expression and promoter accessibility for the overlapped genes identified in (**a**). (**c**) KEGG pathway enrichment analysis of genes with DARs located in their promoter regions. (**d-f**) TF motifs enriched from MSN DEGs. (**g-i**) Predicted motif activity scores for MA0146.2 in oligodendrocytes and astrocytes (**g**), MA0765.2 in microglia (**h**), and MA0599.1 in neuron progenitor cells (**i**). (**j**) Summary of predicted NRF1 and YY2 biding sites identified among DEGs involved in mitochondrial, ribosomal and proteasomal activities. The relative score (Rel Score) reflects the similarity between DNA sequence patterns and reference position-specific scoring matrices (PSSMs) for known motifs; values closer 1 indicate a higher likelihood of true motif occurrence.

whereas *Slc12a2* showed reduced promoter accessibility but increased expression (**Fig. 3b**). KEGG enrichment analysis from genes with DARs in promoters identified signaling pathways involved in cellular metabolism and growth, including insulin, apelin, Hippo, and PI3K-Akt signaling (**Fig. 3c**). These pathways are known regulators of energy production and protein homeostasis. Consistent with this, promoter accessibility changes were observed in key regulatory genes within these pathways, including *Pik3r2*, *Map2k1*, *Gsk3b*, *Ppp2r2d* (**Suppl. Table 3**), suggesting that PCE affects transcriptional regulation via chromatin remodeling of the upstream signaling nodes.

To further characterize regulatory mechanisms, we performed transcriptional factor (TF) motif enrichment analysis on promoter-associated DARs using Signac FindMotifs function and JASPAR2020 database. After stringent filtering (FDR<0.01 and fold-enrichment > 3), seven enriched TF motifs were identified (**Suppl. Fig. 3a-c**). ChromVAR analysis revealed cell-type-specific motif activity patterns. For examples, motifs associated with Zfx (MA0146.2) were enriched in astrocytes and oligodendrocytes, Etv5 (MA0765.2) in microglia, and Kfl5 (MA0599.1) in neuron progenitors (F**ig. 3d-i**), indicating distinct regulatory programs across cell types. Notably, Nrf1 (MA0506.1) motif showed high predicted activity in neuronal populations and was enriched among genes associated with mitochondrial function (**Fig. 3j**). Sequence analysis using TFinder^23^ further demonstrated that multiple PCE-regulated mitochondrial genes including *Ndufb10* (Complex I), *Cox7a2* (Complex IV), and *Atp5mc3* (Complex V) contain high-confident NRF binding sites. Given the established role of Nrf1 in regulating nuclear mitochondrial genes^24–27^, these findings suggest that altered chromatin accessibility at Nrf1 target sites may impair mitochondrial gene expression after PCE (**Suppl. Table** 2, **Fig. 3j**). In addition, Nrf1 binding motifs were also identified in genes involved in translation machinery (**Fig. 3j, Suppl. Table 3**), hinting a potential link between mitochondrial metabolism and protein synthesis. Consistent with this, protomer regions of many ribosomal genes contained high-scoring sequences for Yy2 and SP family TFs (**Fig. 3g, Suppl. Table 3**), which have been implicated in ribosomal gene regulation^28^.

More than half of high-confidence DARs associated sequences matched Klf5 binding motif (MA0599.1). Klf5 is known to regulate cell proliferation and differentiation and to maintain stemness of certain stem cells^29^. High-affinity binding sites were identified in the promoters of genes such as *Elk4* and *Ppp2r2d* (**Suppl. Fig. 3c**). Elk4 acts as a transcription factor, and Ppp2r2d is part of the protein phosphatase 2 (PP2A) family. Both were known to regulate cell cycle and growth^30,31^(**Suppl. Fig. 3a**), which suggests Klf5 is a master regulator in determining MSN cell fate in the NAc. More broadly, many promoters contain sequences resembling multiple TF motifs (**Suppl. Fig. 3b, Suppl. Table 4**), suggesting combinatorial regulation of gene expression in response to PCE. Furthermore, almost all seven TFs with DARs were involved in regulating cell fate, cell proliferation, or mitochondrial biogenesis (**Suppl. Fig. 3c**).

Collectively, these findings indicate that PCE induces selective changes in chromatin accessibility at regulatory regions controlling mitochondrial function, protein synthesis, and cellular signaling, providing a potential epigenetic basis for the observed transcriptional dysregulation.

### Acute THC challenge exacerbates mitochondrial dysfunction in PCE offspring

Given prior evidence that adolescent acute THC exposure worsens PCE-associated phenotypes with male offspring exhibiting heightened vulnerability^32^, we examined its impact on NAc transcriptional profiles specifically in male offspring. Male PCE offspring were subject to a single dosage of acute THC challenge (2.5 mg/kg, s.c.) at P24, and NAc tissue was collected four hours later for snRNA-seq analysis. After quality control and normalization, 44,215 nuclei from control and 26,228 nuclei from THC-treated rats were resolved into 18 unique clusters (**Fig. 4a**), with similar cell type composition and quality metrics across conditions (**Fig. 4b, Suppl. Fig. 4a-c**).

**Fig. 4.**
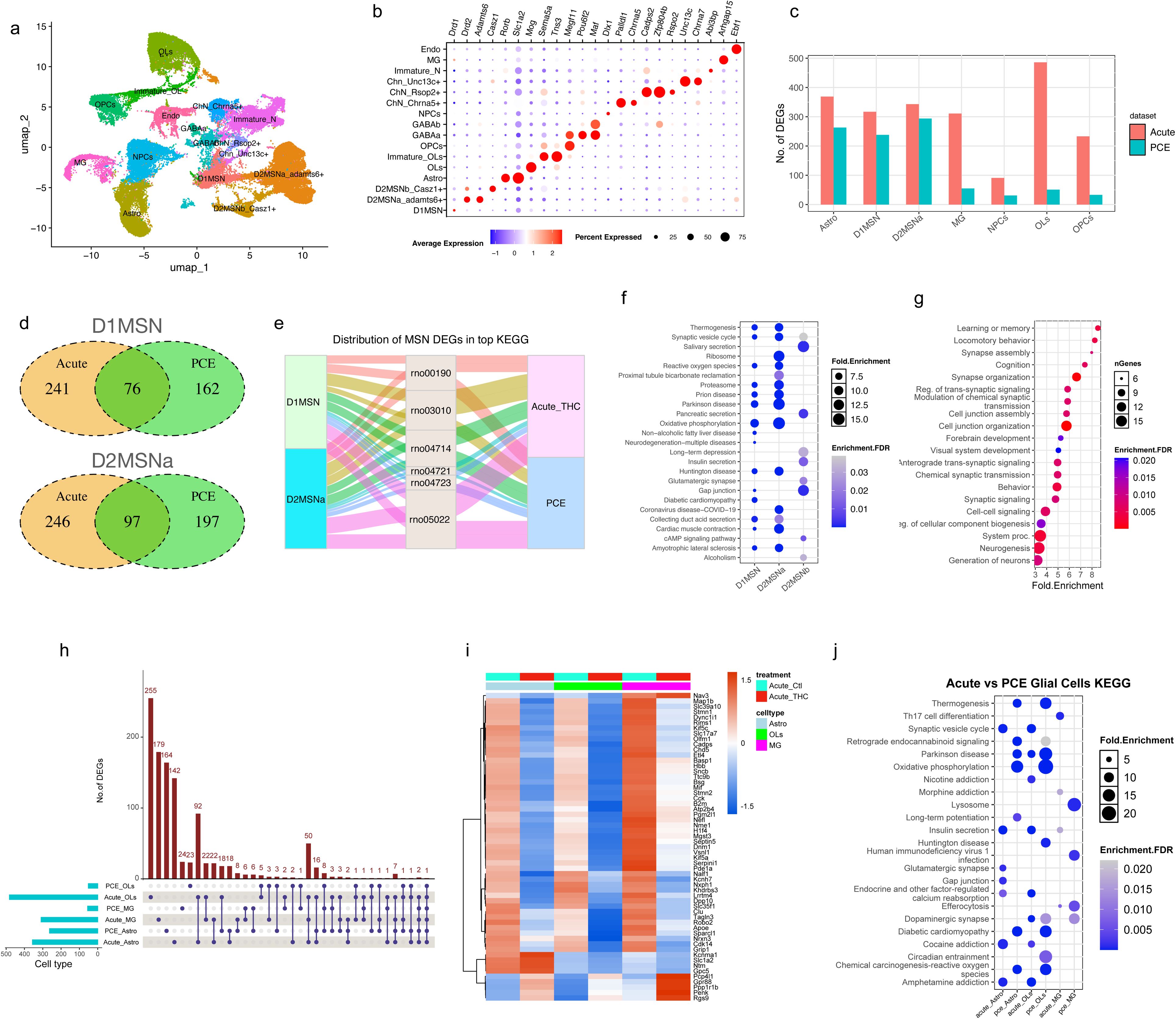
Acute THC challenge exacerbates mitochondrial dysfunction in male PCE offspring nucleus accumbens. (**a**) Unsupervised clustering of single nuclei isolated from the NAc region after acute THC exposure. Each point represents a single nucleus, colored by cell cluster identity. (**b**) Dot plot showing expression of canonical marker genes used to annotate major cell types in the NAc. (**c**) Bar plot showing the number of DEGs in major cell types after PCE alone or acute THC challenge in PCE offspring. Cell types include D1-and D2-medium spiny neurons (D1MSN, D2MSNa, D2MSNb), astro, OLs, MG, and other non-neuronal populations. (**d**) Venn diagrams depicting overlapping DEGs between PCE alone and acute THC challenge in D1MSN and D2MSNa. (**e**) Alluvial plot illustrating the distribution of D1MSNa and D2MSNa DEGs across the top KEGG pathways enriched following PCE alone or acute THC exposure. Pathways shown: oxidative phosphorylation (rno00190); ribosome (rno03010); thermogenesis (rno04714); synaptic vesicle cycle (rno04721); retrograde endocannabinoid signaling (rno04723); neurodegeneration (rno05022). (**f**) Dot plot depicting KEGG pathways enriched from acute THC challenge-induced DEGs in D1 and D2 MSNs. (**g**) Gene Ontology Biological Process (GO: BP) terms enriched from D2MSNb DEGs identified uniquely after acute THC exposure (not present after PCE alone). (**h**) UpSet plot illustrating the number of shared and unique DEGs across major non-neuron cells (astro, OLs, MG) under PCE and acute THC treatments. Horizontal bars show total DEGs per cell type/condition, and vertical bars indicate intersection sizes. (**i**) Heatmap showing expression levels of shared DEGs identified across astro, OLs, and MG following either PCE or acute THC treatment. (**j**) Comparison of KEGG pathways affected by PCE alone versus acute THC exposure in astro, OLs, and MG. Only male P24 rats were selected for acute THC exposure as males were previously found to be more responsive to PCE.

In contrast to PCE, acute THC challenge increased the number of DEGs across major cell types, including D1MSN, MG, OLs, and OPCs (**Fig. 4c, Suppl. Table 5**). Approximately one-third of the PCE-induced DEGs in D1MSN and D2MSNa remained differentially expressed after acute THC challenge (**Fig. 4d**), with similar KEGG pathway enrichment profiles (**Fig. 4e**), indicating reinforcement of PCE-induced transcriptional alteration. The majority of MSNs, particularly D1MSNs and D2-MSNa, exhibited highly concordant responses to acute THC challenge in terms of KEGG enrichment **(Fig. 4f**). Notably, mitochondrial energy production pathways—especially OXPHOS and thermogenesis—were the most enriched. In addition, pathways associated with neurodegeneration diseases, including prion diseases, Parkinson diseases, Huntington diseases, Alzheimer diseases, and amyotrophic lateral sclerosis, were also enriched in MSNs after acute THC administration (**Fig. 4f**). In contrast, DEGs from D2MSNb neurons following acute THC exposure were preferentially enriched in neuronal functions-related pathways, such as synaptic vesicle cycle, gap junction, and long-term depression (**Fig. 4f)**. These pathways were not observed in the PCE condition alone, indicating that acute THC challenge exerts a stronger subtype-specific effects on MSNs. Consistent with this, GO biological process enrichment of D2MSNb-specific DEGs revealed significant associations with learning, memory, cognition, and behavior (**Fig. 4g)**, aligning with the established role of NAc in reward circuitry^12,33^. Other biological processes significantly enriched included synaptic signaling and cell junction organization, and neurogenesis and brain development. Together, these findings indicate that a single acute THC exposure in PCE animals not only amplifies pre-existing transcriptional dysregulation but also induces distinct functional alterations in NAc MSNs.

In glial cells, acute THC exposure markedly increased the number of DEGs (**Fig. 4c**), the majority of which were absent in PCE alone (**Fig. 4h**). A substantial proportion of DEGs were shared across glial cell types: 61 common genes were identified among astrocytes (Astro), oligodendrocytes (OLs), and microglia (MG), and 160 were shared between Astro and OLs after acute THC treatment (**Fig. 4h**). This pattern suggests heightened sensitivity of glial cells to acute THC treatment. Across these cell types, the most shared DEGs were downregulated (**Fig. 4i)** and enriched in synaptic vesicle cycle (**Suppl. Fig. 5a**).

Notably, the expression of *Ppp1r1b* was elevated in the glial cells following acute THC treatment (**Fig. 4i**). In neurons, *Ppp1r1b* encodes a regulatory subunit of protein phosphatase 1 (PP1), which is critical in modulating synaptic plasticity and neuronal signaling^34^. Its upregulation in glia may represent a compensatory response to impaired synaptic vesicle cycling, although its functional role in glial–neuronal interactions remains to be elucidated. Astro and OLs exhibited enrichment of similar KEGG pathways, including synaptic vesicle cycle, calcium signaling, and cocaine- and amphetamine-related pathways (**Fig. 4j**). In contrast, MGs displayed a distinct response profile: acute THC treatment upregulated the transcription of genes associated with immune-related pathways such as Th17 differentiation, B cell signaling, and efferocytosis of the inflammation process, indicating an activation of inflammatory processes in MGs (**Suppl. Fig. 5b-d**).

Together, these results demonstrate that acute THC exposure amplifies PCE-induced transcriptional dysregulation, particularly in mitochondrial and synaptic pathways, while also eliciting additional cell-type-specific responses across neuronal and glial populations.

### THC disrupts mitochondrial transcription which may perturb MSN development and maturation

Mitochondrial function is a critical determinant of neuronal development and maturation^35^ providing both the energy and biosynthetic intermediates required for neurite outgrowth, synaptic formation, and neurotransmission^17,36^. Our transcriptomic and epigenetic analyses indicated that PCE disrupts the expression of OXPHOS genes that may lead to mitochondrial dysfunction. We next investigated whether this impairment affected the development and functional maturation of NAc MSNs.

To this end, we performed Pearson correlation analyses between gene modules of mitochondrial activity and neuronal function. A gene module score for mitochondrial function was calculated by the average expression of genes involved in mitochondrial respiration pathways, including electron transport, OXPHOS, and ATP synthesis. Neuronal function score was similarly defined using genes associated with axon growth, synapse organization, and synaptic signaling.

At P24, both D1 and D2 MSNs in PCE rats exhibited reduced scores for neuronal development and maturation (**Fig. 5a, e**), which were positively correlated with mitochondrial respiration scores in D1 (**Fig. 5b**) and D2 MSNs, respectively (**Fig. 5f**). This correlation suggests that MSN maturation may be delayed due to limited mitochondrial function and metabolic substrates. Acute THC administration further reduced the MSN developmental scores (**Fig. 5i, m**), which was also associated with decreased mitochondrial activity. Consistently, mitochondrial respiration and MSN development showed a moderate but significant positive correlation following acute THC challenge (**Fig. 5j, n**).

**Fig. 5.**
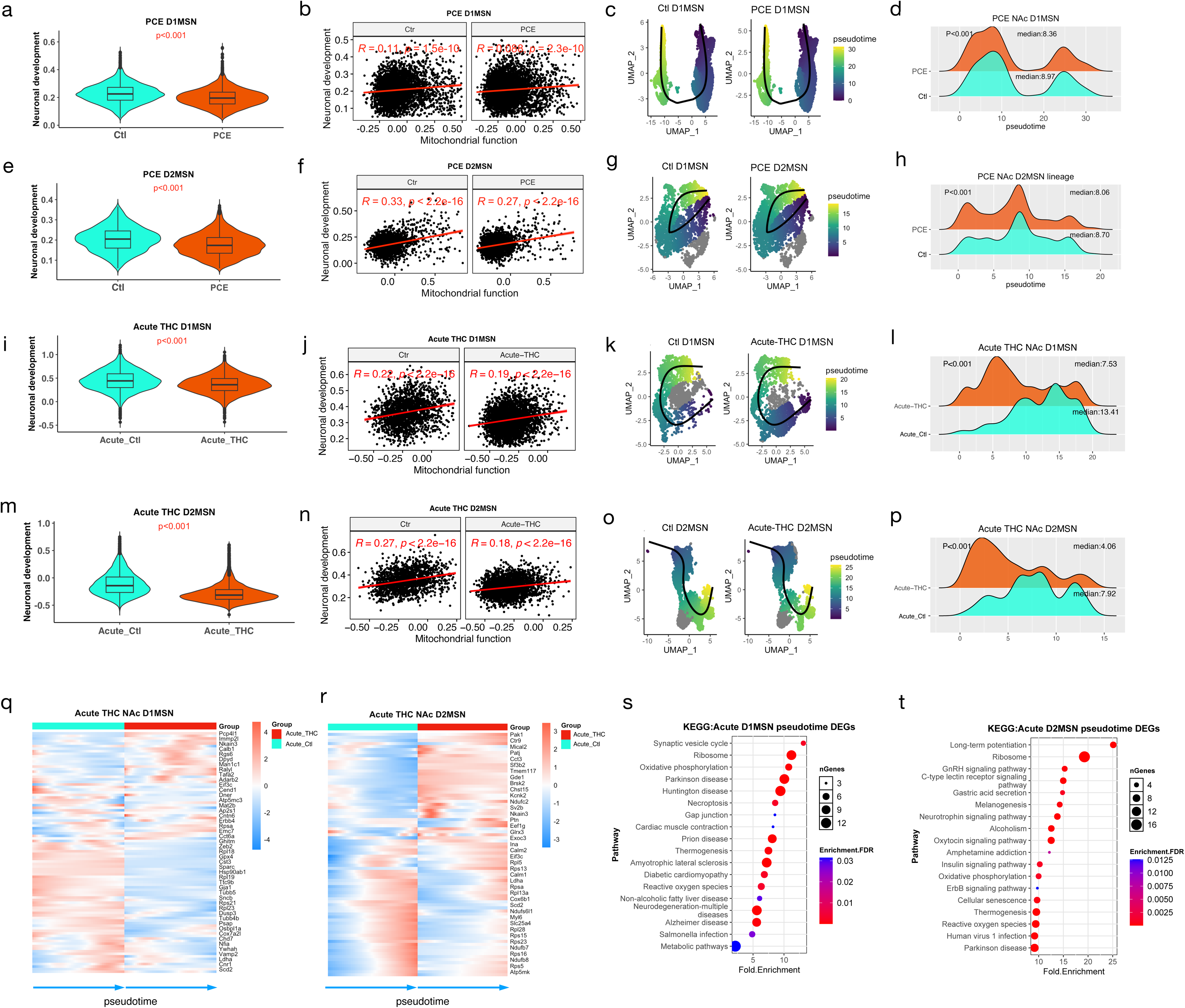
PCE-induced mitochondrial dysfunction is positively associated with neuronal development and maturation. (**a, e, i, m**) Violin plots showing gene expression module scores for neuronal development in PCE D1MSNs (**a**), PCE D2MSNs (**e**), acute THC D1MSNs (**i**), and acute THC D2MSNs (**m**). Box plots indicate the median, 25^th^ percentile, and 75^th^ percentiles. P values were calculated using the Wilcox rank-sum test. (**b, f, j, n**) Pearson correlations between mitochondrial function scores and neuronal development scores in PCE D1MSNs (**b**), PCE D2MSNs (**f**), acute THC D1MSNs (**j**), and acute THC D2MSNs (**n**). Correlation coefficient (R) and p values are shown. (**c, g, k, o**) UMAPs showing inferred trajectories of PCE D1MSNs (**c**), PCE D2MSNs (**g**), acute THC D1MSNs (**k**), and acute THC D2MSNs (**o**). The cells are colored by pseudotime and trajectories (black lines) were inferred using Slingshot. (**d, h, i, p**) Density plots showing the distributions of cells across pseudotime in PCE D1MSNs (**d**), PCE D2MSNs (**h**), acute THC D1MSNs (**i)**, and acute THC D2MSNs (**p**). Median pseudotime values and p values (Wilcox rank-sum test) are indicated. (**q, r**) Heatmaps showing dynamic expression of DEGs induced by acute THC exposure across pseudotime in D1MSNs (**q**) and D2MSNs (**r**). Gene expression trends were fitted using *associationTest* and smoothed using *predictSmooth* function, respectively, from the *tradeSeq* package. (s, t) KEGG pathway enrichment analysis of dynamically regulated acute THC challenge-induced DEGs along the MSN developmental trajectory in D1MSNs (**s**) and D2MSNs (**t**).

To characterize the temporal dynamics of THC-induced transcriptional changes, we performed pseudotime and trajectory analysis of MSN populations. PCE did not alter MSN subcluster composition (e.g., D1 MSNs; **Suppl. Fig. 6a**), or lineage trajectories (**Fig. 5c, g, k, o**). However, pseudotime analysis revealed an increased proportion of MSNs in earlier developmental states in PCE rats (**Fig. 5d, h**), hinting delayed maturation. This delay was exacerbated by an acute THC challenge on P24 PCE offspring (**Fig. 5i, p**). Because lineage trajectories were preserved across PCE and acute THC challenge (**Fig. 5c, g, k, o**), we hypothesized that THC impairs neuronal maturation primarily through transcriptional dysregulation rather than fate specification.

We next examined the dynamics of gene expression along MSN pseudotime. Genes essential for mitochondrial function (e.g., *Cox6b1*, *Cox7a2l*, *Ndufb7)*, and ribosomal activity (e.g., *Rpl28* and *Rpl5)* were activated at later developmental stages and exhibited reduced expression following acute THC challenge (**Fig. 5q, r**). These findings indicate that both reduced energy production and translation capacity may contribute to delayed maturation of MSN. Additionally, genes associated with synaptic activity (*Vamp2* and *Ap2s1)*, and long-term potentiation (e.g., *Calm1* and *Calm2)* were enriched among acute THC-responsive DEGs (**Fig. 5s, t)** and displayed dynamic expression changes along pseudotime (**Fig. 5q, r)**. This is consistent with the prior evidence linking mitochondrial dysfunction to synaptic impairment and neurodegeneration^36^.

To assess cell-type specificity, we randomly selected 2,000 non-neuronal cells from the offspring of PCE and acute THC challenge groups, and subsequently determined their gene module scores and correlations between mitochondrial function and neuronal development. In contrast to MSNs, non-neuronal cells showed no differences in the neuronal development scores and no correlation with mitochondrial functions (**Suppl**. **Fig. 6b-e**), indicating that THC-induced delay in maturation is a neuron-specific effect.

## Discussion

Prenatal cannabis exposure (PCE) has been associated with an increased risk of a wide range of neurological development problems^32,37,38^, however, the mechanism by which PCE affects neuronal development and maturation remain poorly defined. Δ^9^-tetrahydrocannabinol (THC), the primary psychoactive component of cannabis, has been shown to impair mitochondrial energy metabolism by inhibiting oxygen consumption, reducing ATP production, and disrupting mitochondrial membrane potential across multiple cell types^39–41^. However, those observations were largely derived from cytological and pharmacological studies. Here, using single-nucleus transcriptomic and epigenomic approaches, we demonstrated that PCE suppresses gene programs, including components of electron transport chain Complex I, III, and IV, and ATP synthase, consistent with the reduced mitochondrial respiratory capacity observed by other studies^40^.

Notably, the transcriptional dysregulation of nuclear mitochondrial genes was strongly correlated with reduced expression of ribosomal genes (**Fig. 2g, j**), suggesting PCE disrupts transcriptional programs and may alter translational capacity required to maintain mitochondrial homeostasis. This interpretation is consistent with evidence that mitochondrial dysfunction, characterized by increased reactive oxygen species (ROS) formation, altered Krebs cycle metabolism, and reduced ATP production, is tightly coupled to transcriptional regulation and cell fate determination^42,43^. Upstream regulator analysis further identified Zhx2 as a potential regulator of mitochondrial gene expression in PCE offspring, consistent with prior reports that Zhx2 suppresses oxidative phosphorylation by inhibiting transcription of electron transport chain genes and destabilizing PGC-1α^22^.

Sex-specific effects were also observed, with male offspring exhibited a more pronounced reduction in nuclear-encoded mitochondrial gene expression. Male-specific DEGs were enriched in the retrograde endocannabinoid signaling pathways (**Suppl. Fig. 2d**), consistent with evidence that the cannabinoid 1 (CB1) receptor is present at the membrane of neuronal mitochondria (mtCB1), where its activation by exogenous cannabinoids reduces Complex I activity and mitochondrial respiration in neurons^44^ and skeletal muscles^45^. These observation raise the possibility that sex-specific responses may involve differential regulation of cannabinoid signaling pathways that influence mitochondrial gene expression differently between male and female MSNs (**Suppl. Fig. 2g**).

Mitochondrial dynamics and localization are essential for neurite outgrowth, synaptogenesis, and overall neuronal function^17,36,46^. Mitochondria must be trafficked, docked, and maintained at synaptic sites, where they not only provide ATP but also act as scaffolds for the local translation of synaptic machinery assembly^47^. Consistent with this, PCE weakened the correlation between mitochondrial respiratory gene program activity and MSN maturation scores in the NAc, suggesting disruption of the coordinated transcriptional programs that normally couple cellular metabolism to neuronal maturation (**Fig. 5b, f, j, n, s, t**). Motif enrichment analysis revealed significant presence of Nrf1 binding motifs (MA0506.1) in the promoters of PCE-responsive genes (**Fig. 3g**). Nrf1 is a key regulator of nuclear-encoded mitochondrial genes required for respiratory complex assembly^25,27^. In line with this, Nrf1 activity was predicted to be high in MSNs at P24 (**Fig. 3j**), and PCE reduced chromatin accessibility at Nrf1 binding sites, coinciding with the downregulation of mitochondrial genes.

In parallel, motif MA0748.2, recognized by Yy2, was highly active in MSNs (**Fig. 3j**). PCE reduced the expression of ribosomal genes with this motif (**Fig. 3j**), consistent with prior evidence that ribosomal genes are major Yy2 targets^28^. Together, these findings support that coordinated disruption of Nrf- and Yy2-mediated transcriptional programs may contribute to the coordinated suppression of mitochondrial respiratory and protein homeostasis programs.

Importantly, PCE-induced transcriptional changes implicated in multiple neurodegenerative disease, including Parkinson’s disease (PD), Huntington’s disease (HD), amyotrophic lateral sclerosis (ALS), Alzheimer’s disease (AD), and prion disease (**Fig. 2c**, **Fig. 4f, Suppl. Fig. 5s, 5t**). We also observed reduced gene module scores for ribosome and proteasome activity (**Fig. 2f** and **Fig. 6a**), along with the decreased expression of genes encoding these pathways (**Fig. 6b**). The proteasome activity score was positively correlated with mitochondrial function scores, with the strongest correlation observed in D2 MSNs (**Fig 6c**), suggesting that mitochondrial function is crucial for maintaining protein homeostasis.

**Fig. 6.**
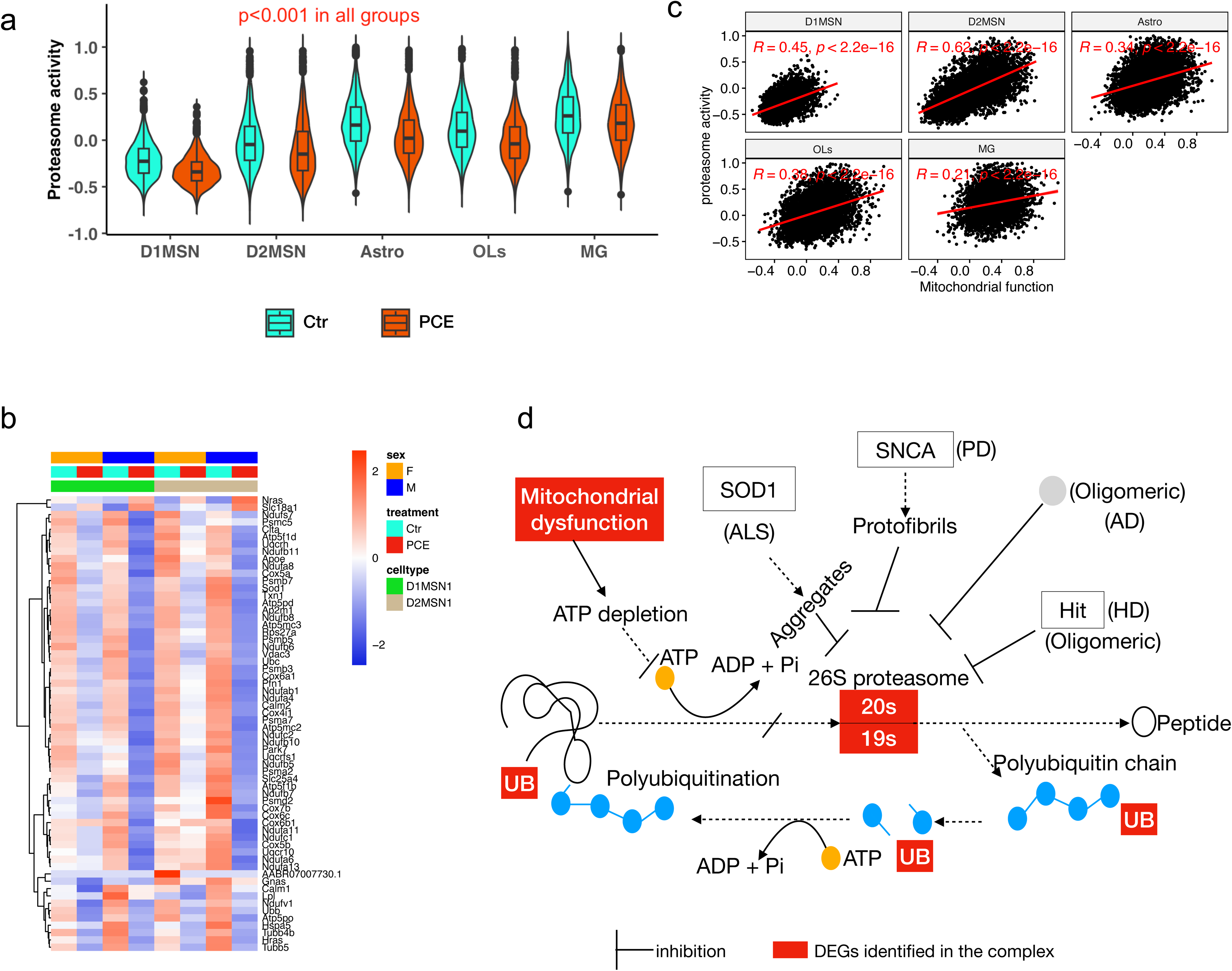
Mitochondrial dysfunction is associated with impaired protein homeostasis and neurodegenerative pathways. (**a**) Violin plots showing gene expression module scores for proteasome activity across five major cell types. (**b**) Heatmap showing the expression of genes involved in protein homeostasis in D1 and D2 MSNs. Normalized snRNA-seq counts were used, and values were scaled by row. (**c**) Pearson correlations between mitochondrial function scores and proteasome activity across five major cell types. (**d**) Schematic illustrating the relationships among mitochondrial dysfunction, proteasome activity, protein homeostasis, and neurodegenerative disease pathways. Unless otherwise indicated, all analyses were performed using combined male and female data from the PCE experiment.

For clarity, we illustrated the potential relationships among mitochondrial respiratory gene disruption, protein homeostasis pathways, and neurodegenerative disease-associated signatures in **Fig. 6d**. In PCE offspring, we speculate that neuron energy production may be restricted due to the dysregulation of mitochondrial gene transcription, which may slow down protein synthesis, reduce ribosome activity and proteasomal degradation, leading to the accumulation of misfolded or damaged proteins which were hallmarks observed in neurodegenerative diseases^48^. Mitochondria are essential for normal neuronal function^17^, neuronal development and synaptic plasticity^49,50^, as well as the regulation of development timing for cortical neurons^17^. Our findings support a hypothesis that dysregulation of mitochondrial metabolic genes following PCE may increase vulnerability to later-life neuronal disfunction.

Integrative transcriptomic and epigenomic analyses revealed Nrf1 and Yy2 as the TFs that coordinately regulate mitochondrial respiration and protein homeostasis. Disruption in these processes after PCE may delay neuronal maturation and exacerbate neurodegenerative processes. Together, this work provides a transcriptional and epigenetic framework linking PCE to altered mitochondrial respiratory gene programs and MSN maturation trajectories in the NAc, highlighting mitochondrial pathways as potential targets for future investigation.

Future studies will be required to determine whether these mitochondrial and proteostatic alterations persist into adulthood and contribute causally to behavioral phenotypes associated with PCE.

## Methods

### Animal care and prenatal THC exposure

All animal experiments involving animal care and sample preparation were approved by the Institutional Animal Care and Use Committee of Loma Linda University. We have complied with all relevant ethical regulations for animal use and the details in animal care were previously described^51^. Pregnant Sprague-Dawley (SD) rats, 3 months old, were purchased from Charles River Laboratories and were randomly divided into two groups: PCE group and control group. THC (20 mg/ml in ethanol) was obtained from the National Institute on Drug Abuse Drug Supply Program, and the THC solution was prepared as described^32^. From gestation day 5 (GD5) to GD20, the animals were administered either THC (2 mg/kg) or vehicle subcutaneously (s.c.) once each day. Offspring was weaned at about postnatal day 21 and maintained without any further manipulation in standard conditions. On P24, eight male rats were divided into two groups and subjected to a one-time acute THC challenge or vehicle (Ctl) s.c. injection (2.5mg/kg). The remaining animals were used for PCE sample collection. We selected P24 as it corresponds to human pre-adolescence which is a sensitive stage for acute THC exposure, and this specific dose has been shown to not induce the typical behavioral responses observed in the cannabinoid tetrad assay, nor does it lead to cannabinoid tolerance^32^. In the acute exposure experiment, samples were collected 4 hours after the s.c. injection.

### Brain tissue dissection

Rat pups on P24 were euthanized by decapitation under deep isoflurane anesthesia. The nuclear accumbens (NAc) regions were isolated on dry ice from the brain slice using a tissue punch (2mm OD). Punctured NAc and mPFC samples were pooled separately from four male or female rats from 3 dams in each group for nuclei isolation and four 10x Multiome single nuclei captures were conducted (M_Ctl, M_PCE, F_Ctl, and F_PCE). We did not use more than two rats of same sex from each litter for the same experiment to control for litter effects. Tissues from acute THC experiment were collected the same way and pooled from four animals from each group for 10x 3’ gene expression study.

### 10x snRNA-seq and Multiome (snRNA-seq and snATAC-seq) library preparation

Single nuclei isolation was performed following the 10x Genomics protocol (CG000124 Isolation of Nuclei for Single Cell RNA Sequencing) and 10,000 nuclei were targeted for each sample in both snRNA-seq and Multiome captures. snRNA-seq libraries were constructed following the established 10x Chromium Next GEM Single-cell 3’ protocol (CG000315 Rev E). For Multiome sample preparation, the resuspended nuclei were subjected to Tn5 transposition proceeding to single cell partitioning into gel beads in emulsion, barcoding. cDNA amplification. The ATAC library construction was conducted by following 10x Genomics protocols CG0001168 Rev C. Library quantification was conducted using Qubit 4.0 (Life Technologies), and quality control was assessed using a TapeStation 2200 and D1000 ScreenTape (Agilent Technologies). All sequencing libraries were prepared at the Center for Genomics at Loma Linda University (LLU). The pooled libraries were sequenced on an Illumina NextSeq 2000 (LLU Center for Genomics) with paired-end sequencing using following setting: Read 1: 28 bp, i7 index: 10 bp, i5 index: 10 bp, Read 2: 90 bp for gene expression libraries and Read 1: 50 bp, i7 index: 8 bp, i5 index: 24 bp, Read 2: 50 bp for snATAC-seq libraries.

### Reads mapping, data quality control, and dimensional reduction

Rnor 7.0 reference genome was downloaded from 10x Genomics (refdata-gex-mRatBN7-2-2024A). snRNA-seq data was mapped and counted using cellranger 8.0.1. Multiome data was mapped using cellrange-arc 2.0.2. Cells with fewer than 500 RNA features and ATAC peaks were removed. The RNA and ATAC matrices were further filtered by removing the 2% cells with the highest reads. The clustering of single nuclei based on RNA profiles was performed using the Seurat package 5.0^52^. Briefly, cell-to-gene counts were normalized, and variable genes were selected for dimension reduction by Principal Component Analysis (PCA), visualized with UMAP and clustered with the Louvain algorithm. snATAC-seq data was analyzed using Signac^53^. Specifically, ATAC common peaks matrices were binarized and normalized by Run Term Frequency Inverse Document Frequency (TF-IDF) method, followed by dimension reduction by PCA, visualization with UMAP on “lsi” reduction. To cluster single nuclei with joint modalities, snRNA-seq data and DNA cell-to-bin data were first subject to dimension reduction using “pca” or “lsi”, respectively. Then, FindMultiModalNeighbors function was used to integrate the two modalities using the first 30 components for “pca” and second to 30th components for “lsi”. Finally, cells were visualized with UMAP using Weighted Nearest Neighbor (WNN) analysis.

### Downstream data analysis

After dimension reduction, distinct clusters were defined by their projections on UMAP and the FindMarkers function was used to extract the cluster markers which were cross-referenced with known cell markers to identify cell types. Differentially expressed genes (DEGs) and differential peaks between treatment (either PCE or acute THC) and control groups were determined using FindMarkers function and significance was defined by “MAST” method with FDR<0.05 and |log2FoldChange| > 0.35. Clusters with fewer than 50 cells were excluded. Differential peaks were annotated to Rnor 7.0 genome using annotatePeak function from R package CHIPseeker^54^. MSN cells were extracted from the PCE or acute THC datasets. Following re-normalization, re-clustering, the UMAP values were used for trajectory analysis using slingshot^55^. Pseudotime values for the major lineage (lineag1) were used for data presentation.

### Canonical pathway and molecular function analysis

Analyses of the gene bio-functional pathways were performed using an online analysis tool ShinyGo (v8.05, http://bioinformatics.sdstate.edu/go/). We also used the IPA (Qiagen) to identify the gene network and upstream regulators. Motif enrichment was performed with the online version of TFinder^56^.

### Statistics and reproducibility

Cell proportion was calculated by combining two subgroups (female and male) in each treatment group (PCE, acute THC challenge and control), respectively. To compare the difference, p values were calculated using Wilcox. For single nucleus RNA-seq and histone data, four animals (from two dams) were pooled in each group (female control, male control, female PCE, and male PCE). The differential expression or differential 5kb bin peak was determined using a non-parametric Wilcoxon rank-sum test as part of the Seurat package. Both p-value and adjusted p-value were reported in single nucleus sequencing data.

### Code availability

The snRNA-seq data were analyzed using Seurat (https://github.com/satijalab/seurat) and the Multiome data was analyzed using Signac (https://github.com/stuart-lab/signac). Slingshot (https://nbisweden.github.io/workshop-archive/workshop-scRNAseq/2020-01-27/labs/compiled/slingshot/slingshot.html) and Tradeseq (https://github.com/statOmics/tradeSeq) was used for trajectory analysis. No in-house tool was used.

## Acknowledgements

The authors would like to thank Ms. Adriana Lopez of the LLU Center for Genomics for her administrative support for the project. The genomic work carried out at the LLU Center for Genomics was funded in part by the National Institutes of Health (NIH) grants S10OD019960 (CW), U01DA058278 (CW), the Ardmore Institute of Health grant 2150141 (CW) and Dr. Charles A. Sims’ gift to LLU Center for Genomics. We gratefully acknowledge the National Institute on Drug Abuse (NIDA) Drug Supply Program for providing the THC used in this study.

## Authors’ contributions

CW conceived, designed the study, and provided funding and resources. ZC, WC, and YS L performed the experiments. ZC analyzed the data, generated all figures and supplementary data, and drafted the manuscript. WC, LG, YD, and JDT revised the manuscript. CW revised and finalized the manuscript. All authors reviewed and agreed with the final version of the manuscript.

## Competing interests

All authors claim there are no conflicts of interest.

## Supplementary Figure Legends

**Suppl. Fig. 1. PCE dataset quality control**

(**a**) UMAP showing the consistency in the single nuclei clustering across the four datasets. (**b**) Violin plots showing the RNA feature and RNA count in each cell type across the four datasets. (**c**) Violin plots showing the ATAC feature and count in each cell type across the four datasets.

**Suppl. Fig. 2. PCE inhibits mitochondrial and ribosomal activity**

Violin plots showing the gene module scores for mitochondrial function in D1MSNa (**a**) and D1MSNa (**b**), and ribosomal activity in D1MSNa (**c**) and D2MSNa (**d**), respectively. (**e**) KEGG enrichment from male specific DEGs induced by PCE in D1MSNa, D1MSNb, and D2MSNa cells. (e) Violin plot showing the expression of mitochondrial CB1 receptor (mtCB1) in all MSN neurons in control animals. Violin plots illustrating the module scores for mitochondrial (**g**) and ribosomal activity (**h**) in astrocytes. (**i)** Pearson correlations between the module scores for ribosomal activity and mitochondrial function in astrocytes within male NAc.

**Suppl. Fig. 3. TF binding motifs enriched based on differential chromatin accessibility peaks induced by PCE in MSNs**

(**a**). Bar plot showing the number of DNA sequences with a Rel. score > 0.9 compared to the known TF motifs. (**b**). UpSet plot illustrating the number of unique and common motif sequences reorganized by the seven TFs identified from a. (**c**). A table summarizing the functions of the TFs and the top potential target sequences enriched from the DARs induced by PCE. Rel. score is a measure of the similarity between a DNA sequence and a known motif sequence. The closes it is to 1, the more likely it is a true TF binding sequence. The motif enrichment was performed using the differential peaks that were annotated to a promoter (TSS±2k bp).

**Suppl. Fig. 4. Acute THC challenge dataset quality control**

(**a**) UMAP showing the cell types identified in NAc after acute THC challenge. Violin plots showing the distributions of RNA feature (**b**) and RNA count (**c**) in each cell type in NAc, respectively.

**Suppl. Fig. 5. Effects of acute THC challenge on glial cells in PCE offspring**

(**a**) KEGG pathway enriched from the common downregulated DEGs in glial cells after acute THC exposure. Bar plots showing the log2FC of DEGs in microglia following an acute THC challenge, related to inflammation responses: Th17 cell differentiation (**b**), B cell signaling (**c**), and efferocytosis (**d**), respectively.

**Suppl. Fig. 6. Randomization control analysis on the effects of acute THC exposure on neurodevelopment in PCE offspring**

(**a**) UMAP showing the similar cluster pattern of D1 MSN subclusters in Ctl and PCE samples. Violin plots showing the gene module score for neurodevelopment in 2000 randomly selected non-neurons in PCE (**b**) and acute THC (**c**) dataset, respectively. Pearson correlation between the neuronal development module score and mitochondrial function score in the random non-neurons in PCE (**d**) and acute THC (**e**).

## Notes

### Competing Interest Statement

The authors have declared no competing interest.

### Summary of Updates

We revised the abstract, results, and discussion sections of the manuscript.

